# A phage-encoded small RNA that mimics chimeric guide to inhibit CRISPR-Cas9

**DOI:** 10.64898/2026.06.03.729762

**Authors:** Ying Chen, Peng Zhao, Chenmin Guo, Dandan Chen, Caihong Yun, Jiyun Chen, Liang Liu

## Abstract

CRISPR-Cas9 relies on a dual crRNA–tracrRNA guide, yet whether RNA-based anti-CRISPRs exist for this system has remained unknown. Here we identify a phage-encoded small non-coding RNA, rAcrIIA1, that adopts an unexpected chimeric crRNA–tracrRNA architecture, faithfully recapitulating the entire single-guide RNA (sgRNA) of Cas9. Cryo-EM structures of the Cas9–rAcrIIA1 binary complex and the Cas9–rAcrIIA1–DNA ternary complex reveal that rAcrIIA1 engages Cas9 nearly identically to the native sgRNA, competitively blocking crRNA and tracrRNA loading and thereby abolishing DNA interference. Strikingly, beyond its natural inhibitory function, rAcrIIA1 is fully reprogrammable: with variable guide sequences, it directs Cas9 for sequence-specific DNA cleavage with efficiency comparable to the canonical sgRNA, and it activates *trans*-cleavage more potently. Thus, rAcrIIA1 provides the first experimentally validated, mechanistically resolved RNA-based inhibitor of CRISPR-Cas9, reveals a distinct anti-CRISPR mechanism through complete sgRNA mimicry, and serves as a versatile bifunctional scaffold for genome editing and diagnostics.

## INTRODUCTION

Bacteria deploy CRISPR–Cas systems as adaptive immune barriers against bacteriophages and other mobile genetic elements^1–8^. These systems incorporate short DNA fragments from invaders into CRISPR arrays and use the transcribed CRISPR RNAs (crRNAs) to guide Cas nucleases toward complementary nucleic-acid targets^5,9,10^. In the type II CRISPR–Cas system, a second RNA component—the trans-activating crRNA (tracrRNA)—is required to process pre-crRNA and to assemble a functional Cas9 effector complex^11–13^. The natural dual-RNA guide can be fused into a chimeric single-guide RNA (sgRNA)^14^, a design that lies at the heart of the now-ubiquitous genome-editing platform^15–18^.

In response, phages have evolved anti-CRISPR (Acr) molecules that neutralize host immunity^19^. Most known Acrs are proteins^20–25^, and several act on the Cas9 system through diverse mechanisms^26–38^: AcrIIA17, for instance, blocks Cas9–sgRNA complex formation, whereas AcrIIA18 induces Cas9-dependent truncation of the sgRNA, rendering it inactive^39,40^. Over the past few years, a different mode of immune evasion has come into view: RNA-based anti-CRISPRs (rAcrs)^41,42^. These small non-coding RNAs mimic the repeat sequences of CRISPR arrays and can displace native crRNAs from their cognate Cas effectors. rAcrs have now been described for type I-C, I-E, I-F, V-A and VI-A systems—all of which rely on a single crRNA guide^41,43,44^. In the type I-F system, a prophage-encoded Racr interacts with Cas6f and Cas7f to stall effector-complex assembly. For type V-A (Cas12a), predicted rAcrs have been identified bioinformatically, and for type VI-A (Cas13), the recently characterized rAcrVIA1 adopts a crRNA-like fold despite negligible sequence similarity, locking the HEPN nuclease in an inactive state. Yet, despite the discovery of rAcr candidates for almost all major CRISPR types, a fundamental gap remains: does an RNA-based inhibitor exist for the type II CRISPR-Cas9 system, with its dual-guide requirement and its unique dependence on both crRNA and tracrRNA?

Here we report the discovery of rAcrIIA1, a phage-encoded small non-coding RNA that disables CRISPR-Cas9 immunity through an unprecedented mechanism. rAcrIIA1 carries 5′ sequence homology to the crRNA repeat and 3′ homology to tracrRNA, thereby adopting a chimeric architecture that remarkably resembles the engineered sgRNA of Cas9. High-resolution cryo-EM structures of the Cas9–rAcrIIA1 binary complex and the Cas9–rAcrIIA1–DNA ternary complex reveal that rAcrIIA1 engages Cas9 in a manner nearly identical to that of the native sgRNA. Functionally, rAcrIIA1 binds directly to Cas9, competes with the native crRNA and tracrRNA, and potently blocks Cas9-mediated DNA interference. Notably, although rAcrIIA1 is genomically adjacent to an AcrIIA18 homologue, the protein shows negligible inhibitory activity because it lacks the conserved catalytic residues required for sgRNA truncation. Beyond its natural role as an immune antagonist, rAcrIIA1 can be reprogrammed with arbitrary guide sequences to direct Cas9 for sequence-specific DNA targeting both *in vitro* and in *in vivo*, with efficiency comparable to that of the canonical sgRNA. This bifunctional nature—an anti-CRISPR RNA that also serves as a programmable guide—establishes rAcrIIA1 as a new molecular tool for genome editing, gene silencing and diagnostics. Our findings close a major gap in the anti-CRISPR landscape, expand the mechanistic repertoire of RNA-based immune evasion to the type II system, and introduce a versatile scaffold for Cas9-based technologies.

## RESULTS

### Phage-derived small RNA rAcrIIA1 blocks CRISPR-Cas9 interference

To search for potential RNA-based inhibitors of the type II CRISPR-Cas9 system, we performed sequence alignment between bacteriophage and bacterial genomes (Figure 1). This analysis revealed a region within the genome of *Streptococcus phage Javan584* (Sj) that exhibited significant homology to the DNA sequences encoding both components of the *Streptococcus pyogenes* (Spy) CRISPR-Cas9 guide apparatus—the crRNA and the tracrRNA (Figure 1A-D). This homologous segment lies within a 545-bp non-coding intergenic region situated between two open reading frames, QBX30964.1 and QBX30963.1 (Figure 1A). Promoter prediction identified canonical –10 and –35 elements at the 5′ end of this region. To determine whether this putative promoter drives transcription, we cloned the entire intergenic fragment into *Escherichia coli* and subjected the expressed RNA to small-RNA sequencing. A single RNA species of approximately 111 nucleotides was specifically transcribed from this locus (Figure 1B). Notably, the 5′ end of this small RNA showed extensive sequence homology to the *S. pyogenes* crRNA (Figure 1C), whereas its 3′ end exhibited strong homology to the tracrRNA (Figure 1D). Given that previously characterized rAcrs for type I, V-A and VI-A systems are defined by their crRNA-mimicry, we hypothesized that this small RNA might act as an RNA-based inhibitor of the type II CRISPR-Cas9 system and designated it rAcrIIA1.

**Figure 1.**
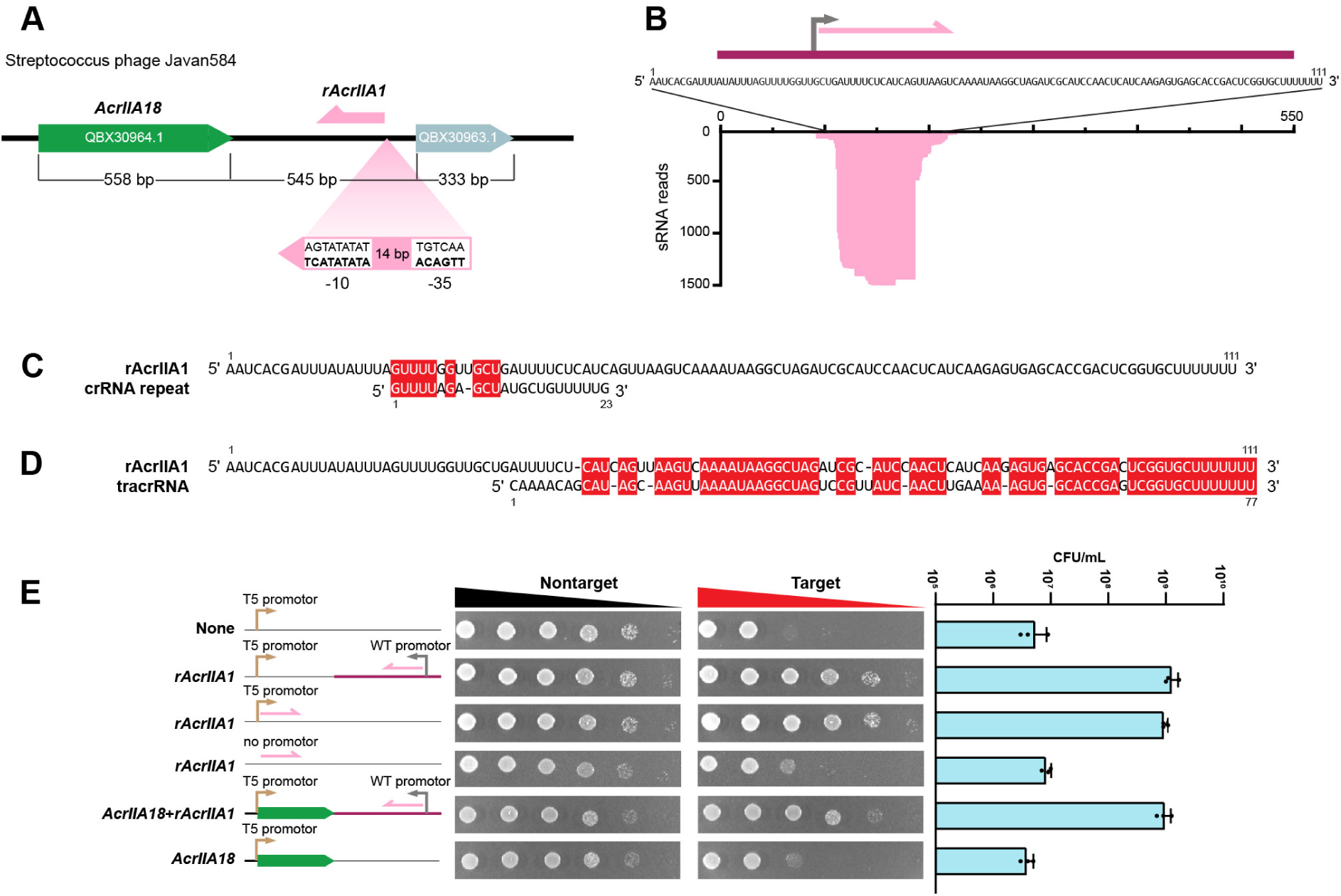
Phage-encoded small RNA rAcrIIA1 blocks CRISPR-Cas9 interference. (**A**) Genomic organization of the *Streptococcus phage Javan584* locus carrying *rAcrIIA1* and *AcrIIA18*. Putative –10 and –35 promoter elements that drive transcription of *rAcrIIA1*. (**B**) Small-RNA sequencing reads aligned to the *rAcrIIA1* locus upon expression of *rAcrIIA1* in *E. coli*. (**C**) RNA sequence alignment of rAcrIIA1 with the *Streptococcus pyogenes* crRNA repeat. (**D**) RNA sequence alignment of rAcrIIA1 with the *Streptococcus pyogenes* tracrRNA. (**E**) SpyCas9-mediated plasmid targeting inhibition assays evaluating the inhibitory ability of rAcrIIA1 or AcrIIA18. crRNA and tracrRNA were used as the guide RNA for SpyCas9 targeting. Gray and yellow arrows indicate the wild-type promoter and the T5 promoter, respectively. Data are represented as mean ± SD (n = 3).

We next tested whether rAcrIIA1 could functionally block CRISPR-Cas9-mediated DNA interference by co-transforming *E. coli* with plasmids carrying rAcrIIA1 and a functional type II CRISPR-Cas9 system (Figure S1A). In the absence of rAcrIIA1, the CRISPR-Cas9 system efficiently targeted a compatible plasmid and severely suppressed bacterial growth. By contrast, co-expression of rAcrIIA1 completely relieved this growth inhibition (Figure 1E and Figure S1B), indicating that rAcrIIA1 potently abrogates CRISPR-Cas9-targeted DNA interference. To determine whether transcriptional expression of rAcrIIA1 is required for its anti-CRISPR function, we compared the activity of rAcrIIA1 driven by its native promoter versus a strong exogenous T5 promoter, alongside a promoter-less control. Both promoter-containing constructs robustly inhibited Cas9-mediated plasmid targeting, whereas the promoter-less rAcrIIA1 failed to do so. These results collectively demonstrate that transcription of rAcrIIA1 is essential for its ability to suppress CRISPR-Cas9 interference.

### rAcrIIA1 outcompetes AcrIIA18 in inhibiting CRISPR-Cas9

In previously characterized type I, V and VI rAcr systems, the *racr* locus often resides adjacent to an anti-CRISPR protein gene^41,44^, with both components capable of independently suppressing CRISPR-Cas immunity. We therefore examined whether rAcrIIA1 is genomically associated with a protein-coding anti-CRISPR gene. The gene immediately downstream of rAcrIIA1 (QBX30964.1) encodes a 185-amino-acid protein that shares 45% sequence identity with the anti-CRISPR protein AcrIIA18 from *Streptococcus macedonicus* (Sm)^39,40^ (Figure 1A and Figure 2A). Structural prediction and comparative modelling further revealed that this protein adopts a three-dimensional fold highly similar to that of AcrIIA18 (Figure 2B and 2C). Based on this strong sequence and structural conservation, we concluded that QBX30964.1 encodes an AcrIIA18 homologue. The upstream gene (QBX30963.1) encodes a 110-amino-acid protein with no detectable homology to any known protein, and its function remains unknown.

**Figure 2.**
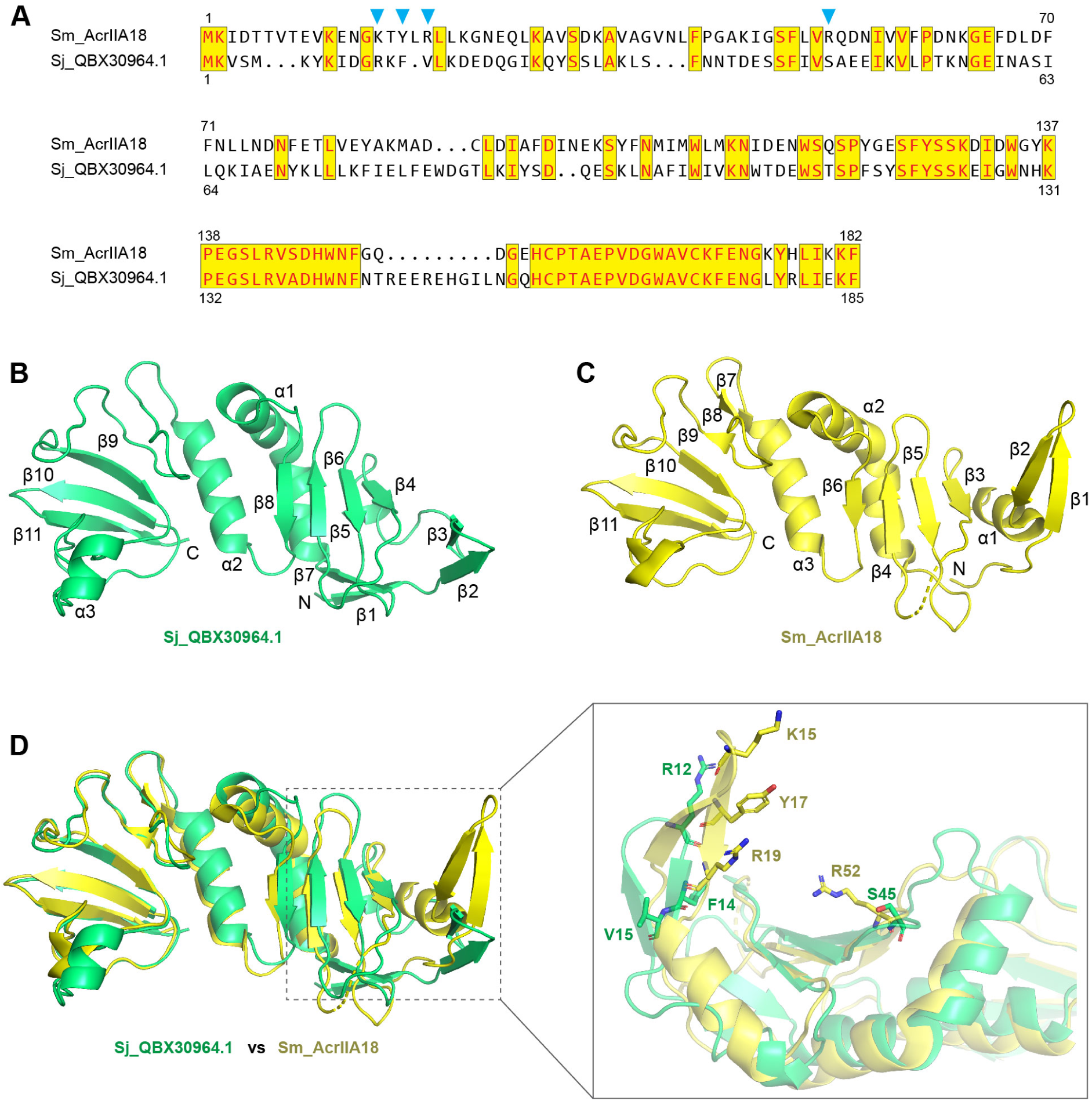
Phage-encoded Sj_AcrIIA18 lacks conserved RNase catalytic residues. (**A**) Sequence alignment of the QBX30964.1 protein from *Streptococcus phage Javan584* (Sj) with AcrIIA18 from *Streptococcus macedonicus* (Sm). Identical residues are shown in red and highlighted in yellow. The RNA catalytic residues of Sm_AcrIIA18 are marked by blue triangles. (**B**) Predicted structure of the Sj_QBX30964.1 protein. (**C**) Structure of Sm_AcrIIA18 (PDB code: 7VLM). (**D**) Structural superposition of Sj_QBX30964.1 (green) and Sm_AcrIIA18 (yellow). The inset provides a magnified comparison of the potential RNase catalytic site, with putative catalytic residues shown as sticks

We next evaluated the inhibitory activity of the phage-derived AcrIIA18 (Sj_AcrIIA18) against CRISPR-Cas9 using bacterial growth inhibition and *in vitro* cleavage assays. Remarkably, in contrast to rAcrIIA1, AcrIIA18 did not exhibit appreciable inhibition of Cas9-targeted DNA interference (Figure 1E and Figure S1B). Previous studies have established that Sm_AcrIIA18 inhibits Cas9 by inducing Cas9-dependent truncation of the sgRNA and that the N-terminal β-hairpin of Sm_AcrIIA18 is critical for this sgRNA-cleavage activity^39^. However, the key catalytic residues within this β-hairpin are not conserved in the phage-derived AcrIIA18 homologue (Figure 2D). Consistent with this observation, our *in vitro* assays failed to detect strong sgRNA degradation activity for the phage AcrIIA18 (Figure S2). Thus, the loss of essential catalytic residues likely accounts for the diminished inhibitory capacity of the phage-encoded AcrIIA18, underscoring that rAcrIIA1—not its protein-coding neighbour—is the primary anti-CRISPR effector at this locus.

### rAcrIIA1 adopts a chimeric crRNA–tracrRNA architecture

Secondary structure prediction of rAcrIIA1 revealed a remarkable architectural resemblance to a chimeric fusion of the crRNA and tracrRNA—the precise configuration of an engineered single-guide RNA (sgRNA) (Figure 3A). Specifically, rAcrIIA1 contains both the crRNA–tracrRNA duplex region and the three stem-loop motifs characteristic of the tracrRNA. To map the functional determinants of rAcrIIA1, we generated a series of deletion mutants and tested their ability to block CRISPR-Cas9 interference. Deletion of sequences immediately downstream of rAcrIIA1 (designated seq1 and seq2) did not affect inhibitory activity, whereas truncations within the rAcrIIA1 core region substantially compromised function (Figure 3B). Both the crRNA–tracrRNA duplex and the three tracrRNA stem-loops were required for full activity; however, the stem-loop deletions imposed the most severe loss of inhibition. These findings demonstrate that rAcrIIA1 suppresses CRISPR-Cas9 by adopting a complete sgRNA-mimetic architecture, and that preserving the integrity of this chimeric crRNA–tracrRNA structure is essential for its anti-CRISPR function.

**Figure 3.**
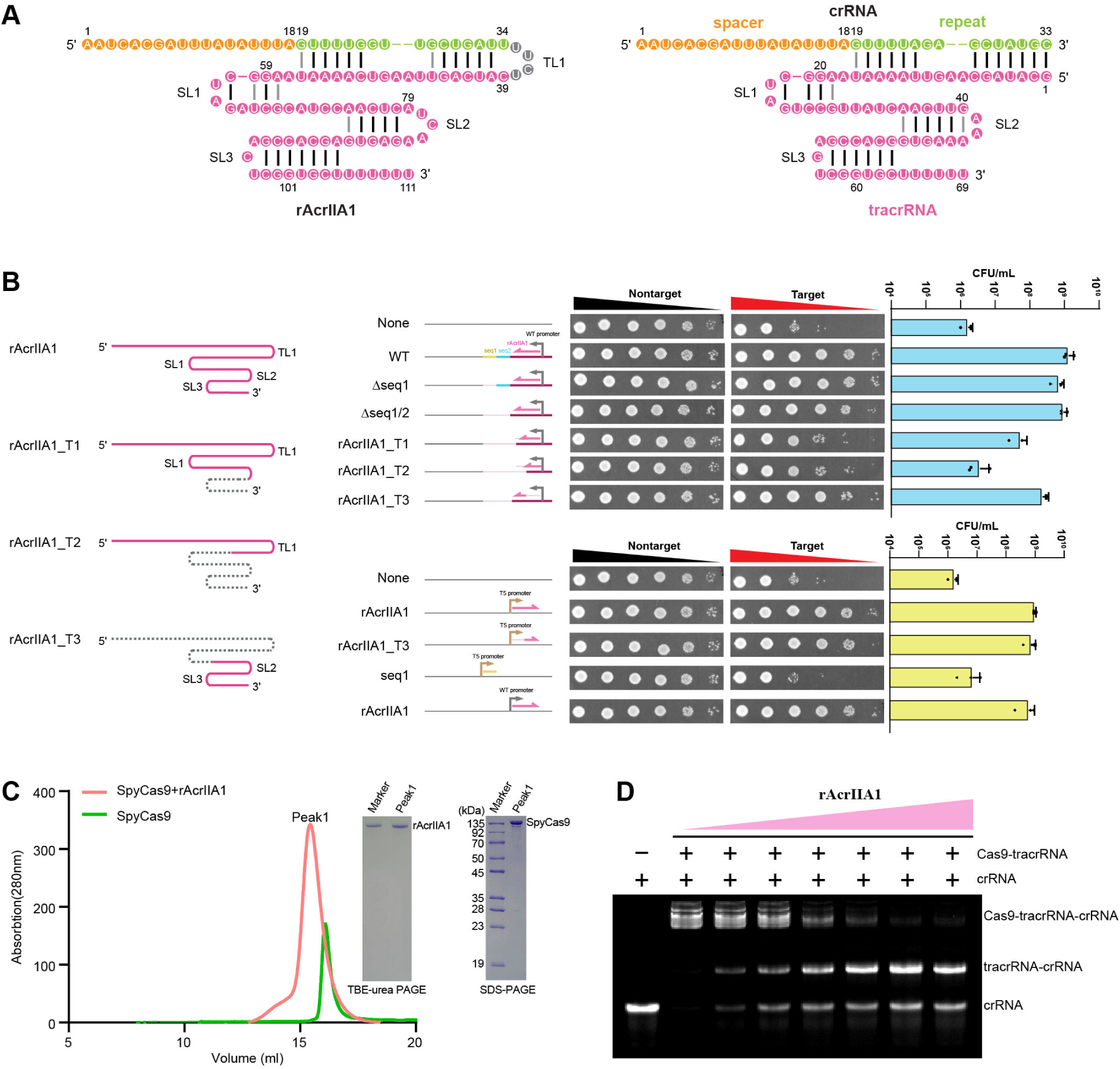
rAcrIIA1 mimics a chimeric crRNA–tracrRNA architecture to inhibit Cas9. (**A**) Secondary structure diagrams of rAcrIIA1 (left panel) and the crRNA–tracrRNA duplex (right panel). The tracrRNA is highlighted in pink; the crRNA repeat and spacer are highlighted in green and yellow, respectively. (**B**) Effects of rAcrIIA1 truncations on the ability to inhibit SpyCas9-mediated plasmid targeting. Data are represented as mean ± SD (n = 3). (**C**) Size exclusion chromatography (SEC) analysis of rAcrIIA1 binding to SpyCas9. SDS-PAGE and TBE-urea PAGE showing that SpyCas9 and rAcrIIA1 co-elute in the SEC fraction. (**D**) Electrophoretic mobility shift assay showing that rAcrIIA1 competes with crRNA and tracrRNA for binding to SpyCas9. The crRNA was 5′-end labeled with FAM.

### rAcrIIA1 directly and tightly binds Cas9

Given that rAcrIIA1 structurally mimics an sgRNA, we hypothesized that it might directly interact with Cas9. Size-exclusion chromatography (SEC) confirmed that purified rAcrIIA1 binds directly to Cas9 *in vitro*, forming a stable protein–RNA binary complex (Figure 3C). To test whether this interaction occurs in the cellular context, we co-expressed rAcrIIA1 and Cas9 in *E. coli*, purified the complex using affinity and ion-exchange chromatography, and analysed the eluate by electrophoresis and small-RNA sequencing (Figure S3). Both approaches unequivocally demonstrated the presence of rAcrIIA1 in the purified Cas9 complex, confirming that rAcrIIA1 associates with Cas9 inside bacterial cells.

We next asked whether rAcrIIA1 competes with the native crRNA and tracrRNA for Cas9 binding. Electrophoretic mobility shift assays (EMSA) showed that increasing concentrations of rAcrIIA1 progressively displaced the crRNA and tracrRNA from pre-assembled Cas9 complexes, indicating that rAcrIIA1 outcompetes the native guides for Cas9 binding (Figure 3D). Consistent with this competitive inhibition model, rAcrIIA1 binding precluded the loading of functional crRNA–tracrRNA into Cas9, thereby blocking guide-directed, sequence-specific DNA targeting. These results establish that rAcrIIA1 functions as a competitive RNA inhibitor that binds directly to Cas9 and prevents assembly of the native CRISPR-Cas9 interference complex.

### Structure of Cas9 in complex with rAcrIIA1

To elucidate the molecular basis of CRISPR-Cas9 inhibition by rAcrIIA1, we determined the cryo-electron microscopy (cryo-EM) structure of the Cas9–rAcrIIA1 binary complex at 3.03 Å resolution (Figure 4, Figure S4, and Table S1). In this binary complex, Cas9 adopts a bilobed architecture resembling that observed in the Cas9–sgRNA complex (Figure 4A and 4B)^45^, comprising the recognition (REC) lobe and the nuclease (NUC) lobe^46^. The REC lobe consists of three α-helical domains (REC1, REC2 and REC3), whereas the NUC lobe comprises the HNH, RuvC, WED, PI and bridge helix (BH) domains. The BH domain, formed by a long α-helix, connects the REC and NUC lobes. A large groove between the two lobes accommodates rAcrIIA1.

**Figure 4.**
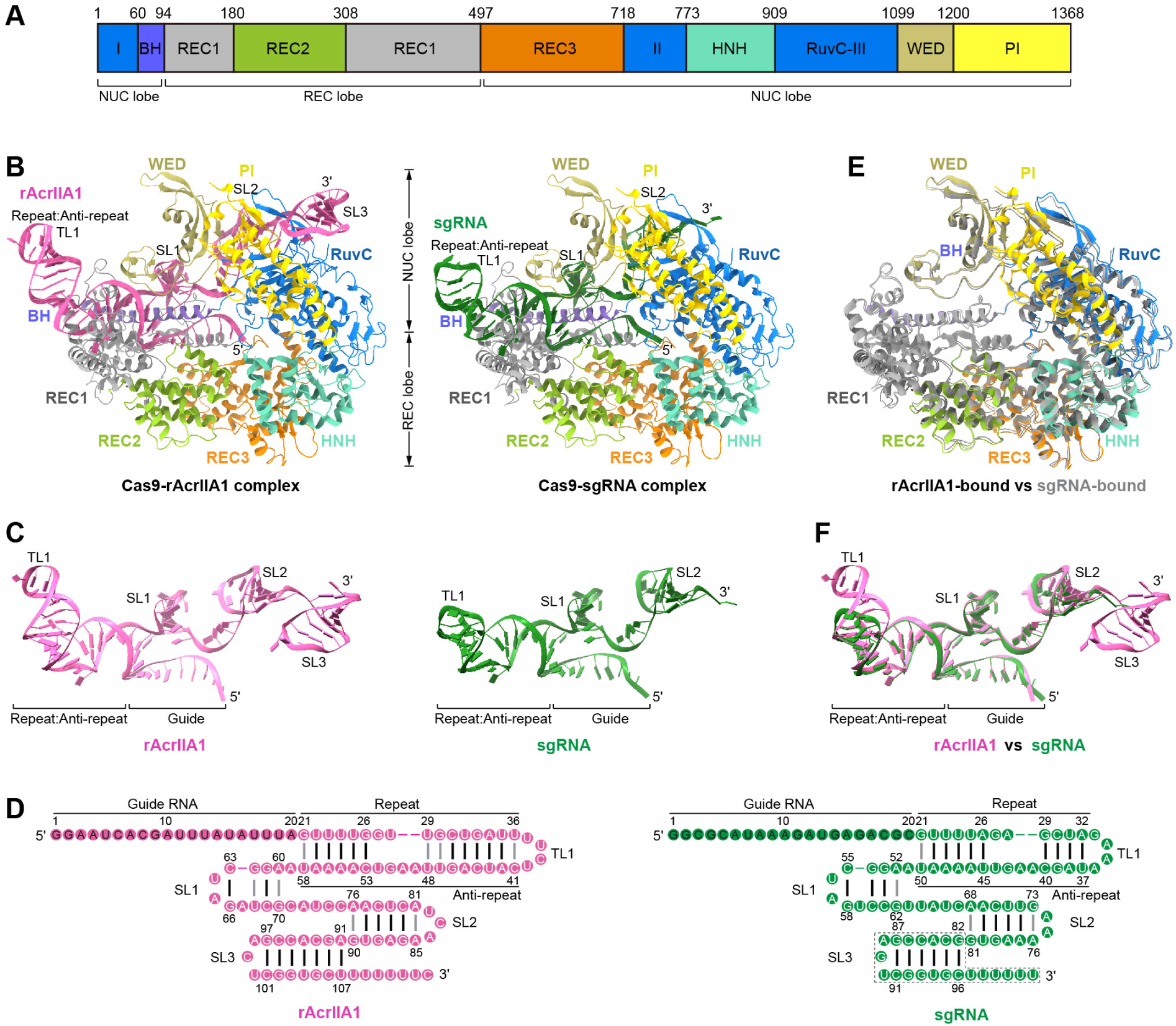
Overall structure of the SpyCas9-rAcrIIA1 binary complex. (**A**) Schematic of domain organization of SpyCas9. (**B**) Overall structures of the SpyCas9-rAcrIIA1 (left) and SpyCas9-sgRNA (right, PDB code: 4ZT0) complexes in cartoon representation. The color-coding used for SpyCas9 is identical to that used in (A). (**C**) The structures of rAcrIIA1 (left) and sgRNA (right, PDB code: 4ZT0). The rAcrIIA1 and sgRNA are shown in pink and green, respectively. (**D**) The schematic of the rAcrIIA1 (left) and sgRNA (right). (**E**) Structural superposition of SpyCas9 in the rAcrIIA1-bound complex (colored as in (A)) and the sgRNA-bound complex (gray). (**F**) Structural superposition of rAcrIIA1 (pink) from the SpyCas9-rAcrIIA1 complex and sgRNA (green) in the SpyCas9-sgRNA complex.

Strikingly, the overall structure of rAcrIIA1 is highly similar to that of sgRNA in the Cas9–sgRNA complex (Figure 4C and 4D). The 5′ end region of rAcrIIA1 (nucleotides 1–20) resembles the guide region of crRNA. Nucleotides 21–36 correspond to the crRNA repeat region, and nucleotides 41–58 resemble the tracrRNA anti-repeat region (Figure 4C and 4D). The repeat and anti-repeat regions form a 14-base-pair duplex. A tetraloop (nucleotides 37–40) connects the repeat and anti-repeat sequences. Downstream of the anti-repeat, nucleotides 60–70 fold into stem-loop 1 (four base pairs), and the 3′ tail region (nucleotides 91–107) forms stem-loop 3 (seven base pairs). Stem-loop 2 (six base pairs) lies between stem-loops 1 and 3. The 5′ guide region is positioned within the central groove formed by the REC and NUC lobes (Figure 4B). The repeat–anti-repeat duplex is anchored by the REC1 and BH domains, stem-loop 1 is engaged by REC1 and WED, and stem-loops 2 and 3 are recognized and stabilized by the WED, RuvC and PI domains.

Superposition of the Cas9–rAcrIIA1 and Cas9–sgRNA (PDB 4ZT0) complexes revealed that the Cas9 structures align closely (root mean square deviation [RMSD] of 1.26 Å over 1,331 equivalent Cα atoms)^45^, indicating that Cas9 adopts a similar conformation in both complexes (Figure 4E). The RNA architectures also superimpose well, with minor conformational differences localized to the repeat–anti-repeat duplex and stem-loop 3 (Figure 4F). In the duplex, nucleotide U29 forms a non-canonical base pair with U48 in rAcrIIA1 (Figure 4D), which is absent in sgRNA. Stem-loop 3 of rAcrIIA1 contains seven base pairs, whereas sgRNA contains six, with an additional Watson-Crick pair formed by A91 and U107 (Figure 4D). In addition, stem-loop 2 of rAcrIIA1 has a three-nucleotide loop, whereas sgRNA has a two-nucleotide loop. Despite these modest differences, the global architecture of rAcrIIA1 is remarkably concordant with that of sgRNA, reinforcing the conclusion that rAcrIIA1 functions as a structural mimic of the Cas9 guide.

### Structure of Cas9 bound to rAcrIIA1 and target DNA

We next investigated whether rAcrIIA1, like an sgRNA, can direct Cas9 to bind target DNA. Size-exclusion chromatography (SEC) showed that the Cas9–rAcrIIA1 binary complex binds with high affinity to a double-stranded DNA oligonucleotide containing a sequence complementary to the guide region of rAcrIIA1 and a canonical 5′-AGG-3′ PAM motif, forming a homogeneous ternary complex (Figure S5A). To visualize the structural basis of DNA recognition, we determined the cryo-EM structure of the Cas9–rAcrIIA1–DNA ternary complex at 2.78 Å resolution (Figure 5, Figures S5 and S6, and Table S1). For this analysis, we introduced alanine substitutions at two catalytic residues—Asp10 in the RuvC domain and His840 in the HNH domain—to prevent DNA cleavage. The HNH domain was not visible in the electron density map (Figure 5A), indicating an intrinsically flexible conformation, consistent with previous observations in multiple Cas9–sgRNA–DNA ternary complex structures^47,48^.

**Figure 5.**
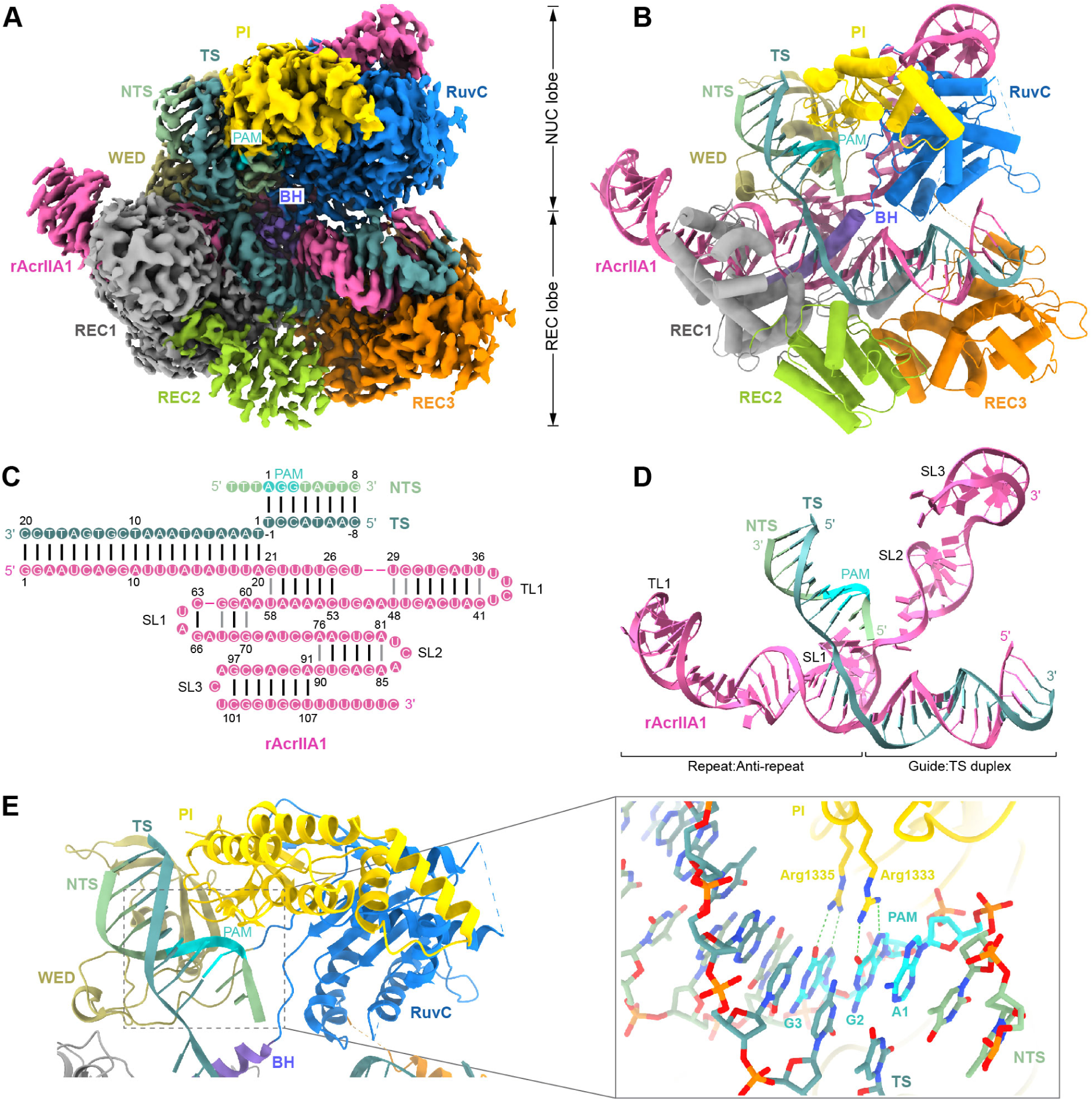
Overall structure of the SpyCas9-rAcrIIA1-DNA ternary complex. (**A**) Cryo-EM density map of SpyCas9-rAcrIIA1-DNA ternary complex. The map is colored by the model. (**B**) Overall structures of the SpyCas9-rAcrIIA1-DNA ternary complex in cartoon representation. (**C**) The schematic of the rAcrIIA1 and target DNA. (**D**) The structures of rAcrIIA1 and target DNA. (**E**) Overall view of the PAM motif bound to the PI domain of SpyCas9, and a detailed view of the interactions between PI-domain residues Arg1333 and Arg1335 and the PAM motif.

In the ternary complex, Cas9 maintains a bilobed architecture similar to that observed in the Cas9–sgRNA–DNA complex (PDB 6O0X) (Figure 5B)^48^. rAcrIIA1 and the target DNA form a structure analogous to the sgRNA–DNA complex (Figure 5C and 5D). The guide sequence of rAcrIIA1 (nucleotides 1–20) hybridizes with the complementary strand of DNA to form a 20-base-pair RNA–DNA heteroduplex (Figure 5C), which resides within the central channel formed by the REC and NUC lobes (Figure 5A and 5B). The DNA duplex region containing the PAM motif is bound in the groove between the PI and WED domains (Figure 5E). Analysis of protein–PAM interactions revealed that the side chains of two arginine residues in the PI domain (Arg1333 and Arg1335) form hydrogen bonds with the G2 and G3 bases of the 5′-AGG-3′ PAM motif, respectively, establishing a PAM recognition mechanism identical to that of sgRNA-guided Cas9.

Superposition of the Cas9–rAcrIIA1–DNA and Cas9–sgRNA–DNA complexes showed that the Cas9 structures align very closely (RMSD of 0.81 Å over 1,102 equivalent Cα atoms) (Figure S5B), and the RNA–DNA heteroduplexes also superimpose with high concordance (Figure S5C). These structural findings demonstrate that rAcrIIA1 can faithfully guide Cas9 to sequence-specifically recognize target DNA, further confirming that rAcrIIA1 is a bona fide structural and functional mimic of sgRNA.

### Reprogrammed rAcrIIA1 directs Cas9 for effective DNA interference

Having established that rAcrIIA1 structurally mimics an sgRNA and binds Cas9 in competition with native guides, we next asked whether rAcrIIA1 could itself direct Cas9 to cleave specific DNA targets. We designed a target DNA sequence complementary to the putative guide region of rAcrIIA1, cloned it into a plasmid, and performed *in vitro* cleavage assays. Robust linearization of the target plasmid was observed in the presence of SpyCas9 and rAcrIIA1 (Figure 6A and Figure S7A). In contrast, no cleavage was detected when rAcrIIA1 was combined with *Staphylococcus aureus* Cas9 (SauCas9) or *Francisella novicida* Cas9 (FnCas9) (Figure S7B and S7C), demonstrating that rAcrIIA1 is a specific guide for SpyCas9. To test whether rAcrIIA1 can be reprogrammed to target arbitrary sequences, we designed a series of rAcrIIA1 variants with different spacer sequences complementary to distinct regions of a plasmid. Each reprogrammed rAcrIIA1 efficiently directed SpyCas9 to cleave its intended target *in vitro* (Figure 6B), confirming that rAcrIIA1 can be engineered as a programmable guide RNA for sequence-specific DNA cleavage.

**Figure 6.**
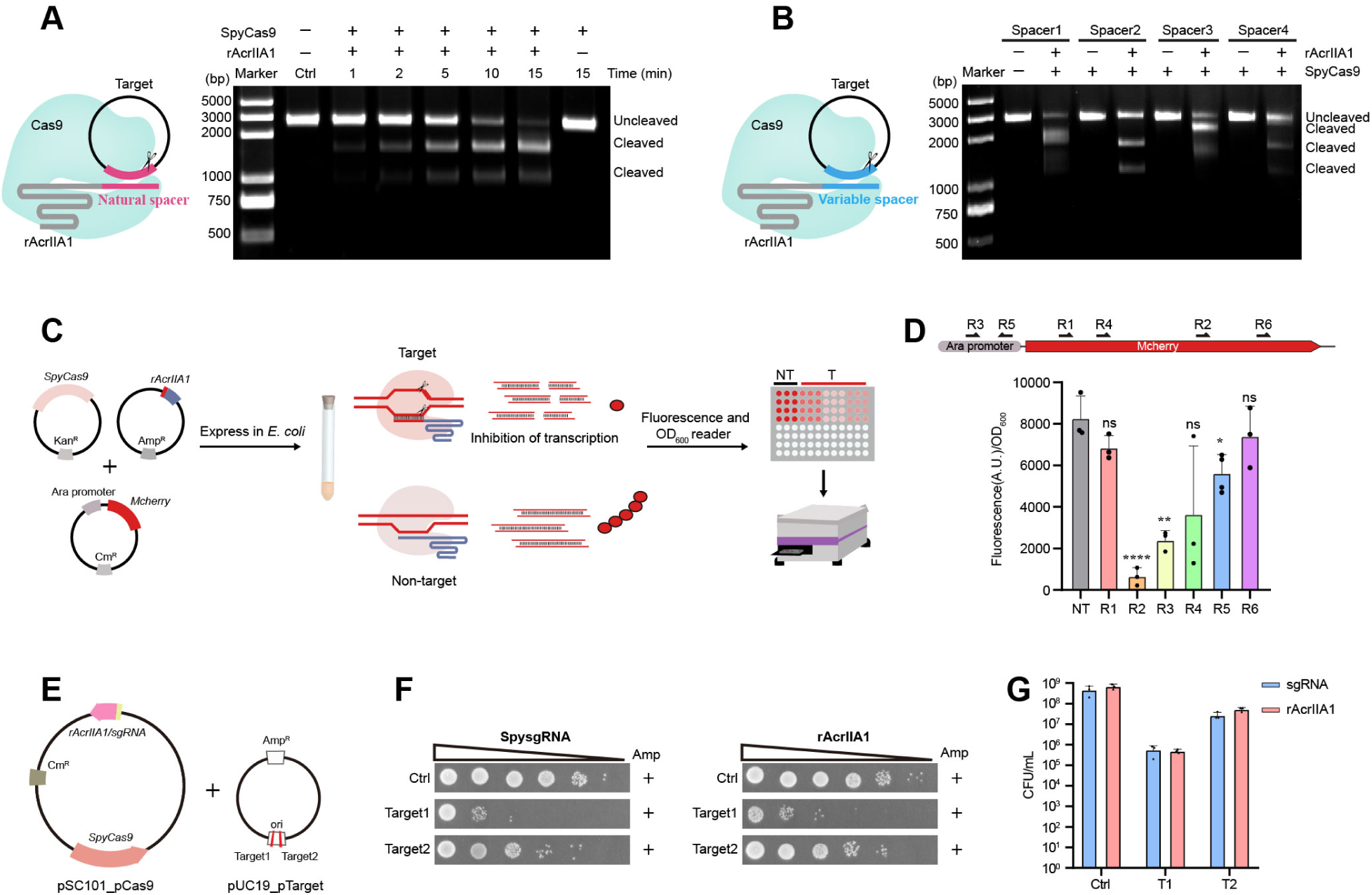
Reprogrammed rAcrIIA1 directs SpyCas9 for effective DNA interference. (**A**) *In vitro* cleavage assays showing rAcrIIA1-mediated SpyCas9 targeting of linear plasmid substrates. (**B**) *In vitro* cleavage assays showing rAcrIIA1 variants with different spacers guiding SpyCas9 to cleave linear plasmid substrates. (**C**) Experimental workflow for the SpyCas9-mediated mCherry fluorescence interference assay in *E*. *coli* using reprogrammed rAcrIIA1 as the guide RNA. (**D**) SpyCas9-mediated silencing of mCherry expression using reprogrammed rAcrIIA1 as the guide RNA. A schematic of the DNA region targeted by different spacers (R1–R6) is shown above. Data are represented as mean ± SD (n = 3). (**E**) Schematic of the rAcrIIA1- or sgRNA-encoding SpyCas9 plasmid and the pTarget plasmid used for DNA targeting and cleavage in *E*. *coli*. (**F**) Growth assays of *E*. *coli* on ampicillin-containing plates, evaluating the ability of reprogrammed rAcrIIA1 or sgRNA to guide Cas9 for targeting and cleavage of an ampicillin-resistance plasmid. (**G**) Colony-forming unit (CFU) comparison for SpyCas9-mediated DNA targeting and cleavage guided by rAcrIIA1 or sgRNA. Data are represented as mean ± SD (n = 3).

Beyond double-stranded DNA cleavage, we examined whether rAcrIIA1 could activate the *trans*-cleavage activity of SpyCas9, a property exploited in molecular diagnostics. Using ssDNA reporters, we found that rAcrIIA1 potently stimulated SpyCas9 *trans*-cleavage (Figure S7D), and notably, it outperformed the canonical sgRNA under the same conditions. This observation suggests that rAcrIIA1-based Cas9 activation could serve as a sensitive platform for nucleic acid detection.

We next investigated whether rAcrIIA1 can mediate Cas9-based gene silencing inside living cells. We designed five rAcrIIA1 guide RNAs targeting either the coding sequence or the promoter region of the *mCherry* reporter gene (Figure 6C). Co-expression of SpyCas9 and rAcrIIA1 targeting the coding region (guide RNA 2) or the promoter region (guide RNA 3) resulted in a marked reduction of mCherry fluorescence (Figure 6D), whereas guides with non-matched sequences had no effect. Thus, rAcrIIA1 can be harnessed for intracellular gene silencing.

To evaluate the specificity and potency of rAcrIIA1 in a physiological context, we engineered an *E. coli* plasmid interference assay (Figure 6E). A guide RNA derived from rAcrIIA1 was designed to target the origin of replication of a compatible plasmid, whose propagation is essential for bacterial growth. When SpyCas9 and the targeting rAcrIIA1 guide were co-expressed, plasmid replication was severely impaired, leading to growth inhibition (Figure 6F and 6G, and Figure 7A). By contrast, expression of SauCas9 or FnCas9 together with the rAcrIIA1 guide did not affect plasmid propagation (Figure 7B and 7C), underscoring the specificity of rAcrIIA1 for SpyCas9. Strikingly, when we compared the activity of rAcrIIA1 with that of the canonical sgRNA in this cellular assay, both guides similarly inhibited plasmid maintenance (Figure 6F and 6G), indicating that rAcrIIA1 and sgRNA possess comparable potency for SpyCas9-mediated DNA targeting *in vivo*. Collectively, these results establish that rAcrIIA1 is not only a natural anti-CRISPR RNA but also a fully programmable and efficient guide RNA that can be engineered for diverse biotechnological applications, including genome editing, gene silencing and molecular diagnostics.

**Figure 7.**
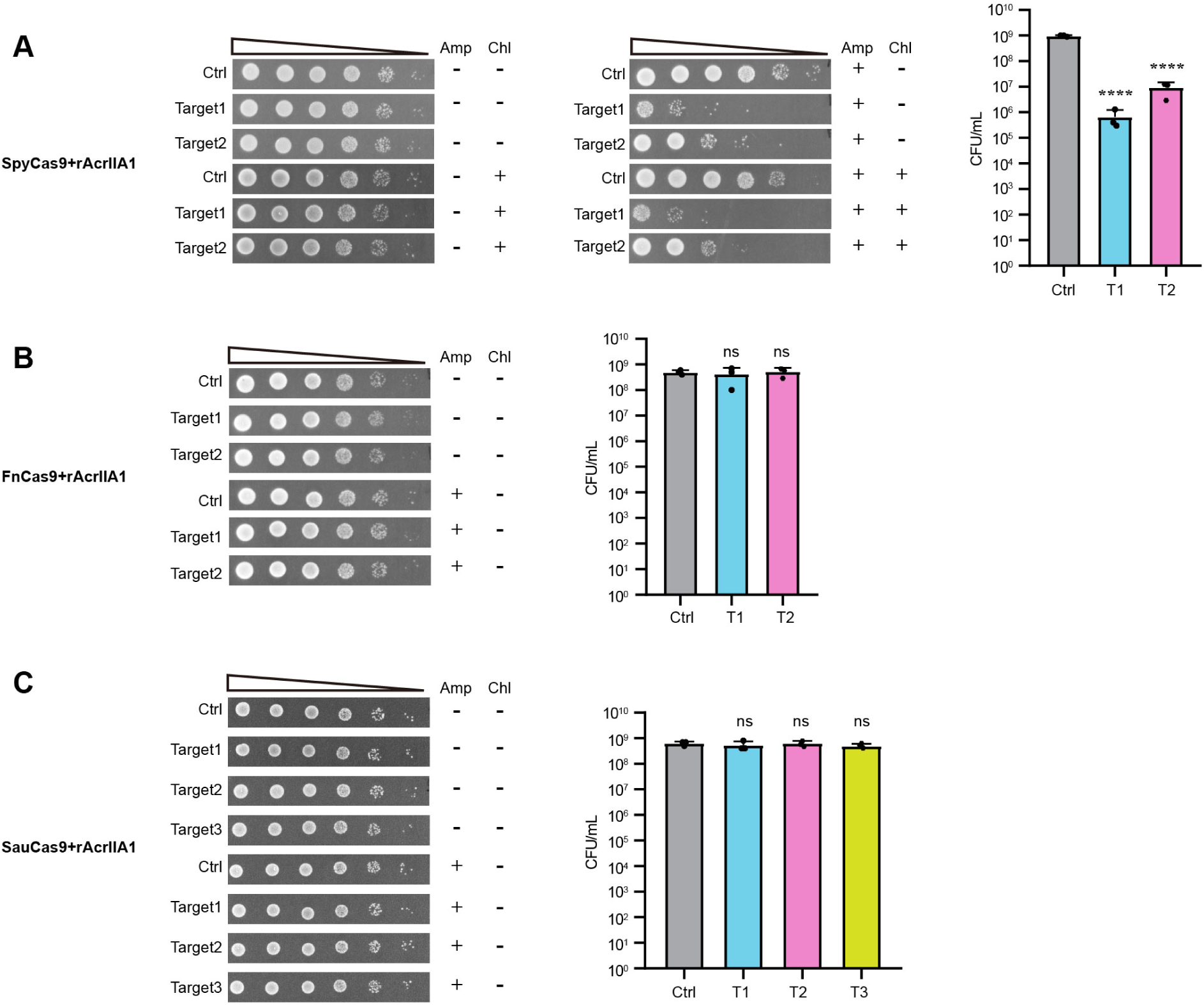
Reprogrammed rAcrIIA1 directs SpyCas9 for specific *in vivo* DNA targeting. (**A**–**C**) Growth assays of *E*. *coli* on ampicillin-containing plates, evaluating the ability of reprogrammed rAcrIIA1 to guide SpyCas9 (A), FnCas9 (B), and SauCas9 (C) for targeting and cleavage of an ampicillin-resistance plasmid.

## DISCUSSION

The discovery of anti-CRISPR RNAs has established RNA-mediated molecular mimicry as a widespread strategy to disarm CRISPR-Cas immunity. Here we identify rAcrIIA1, the first small non-coding RNA inhibitor of the type II CRISPR-Cas9 system, extending the rAcr paradigm from Class 1 (type I-C, I-E, I-F) and Class 2 (type V-A, VI-A) systems to the widely used Cas9. A previous study described a putative Cas9-targeting mechanism involving a phage-encoded, catalytically impaired Cas9 together with a phage-derived tracrRNA, which was proposed to sequester host crRNAs and potentially interfere with host Cas9 function^49^. In contrast, rAcrIIA1 is a single, protein-independent RNA that directly and potently disables Cas9, providing the first experimentally validated example of a pure RNA-based inhibitor for the type II system.

Although all rAcrs exploit mimicry of CRISPR RNA components, their mechanisms diverge in ways that reflect the architecture of the target Cas effector. Type I-F rAcr binds Cas6f and Cas7f to form aberrant subcomplexes^44^, whereas type VI-A rAcrVIA1 locks Cas13 in an inactive HEPN conformation without target RNA engagement^43^. By contrast, rAcrIIA1 adopts a chimeric crRNA–tracrRNA architecture that recapitulates the entire single-guide RNA of Cas9 (Figure 3). Our cryo-EM structures of the Cas9–rAcrIIA1 binary and Cas9–rAcrIIA1–DNA ternary complexes reveal that rAcrIIA1 mimics not only the crRNA guide but also the tracrRNA stem-loops (Figures 4 and 5), achieving near-identical binding to Cas9 as the native sgRNA. This dual mimicry is a unique adaptation to the dual-RNA guide requirement of Cas9, demonstrating that phages can evolve RNA-based inhibitors tailored to the specific guide architecture of each CRISPR type.

An unexpected finding is the functional divergence between rAcrIIA1 and its genomically adjacent protein-coding gene, which encodes an AcrIIA18 homologue. Canonical AcrIIA18 inhibits Cas9 by inducing sgRNA truncation through a conserved N-terminal β-hairpin^39^. However, the phage-derived AcrIIA18 lacks the catalytic residues in this hairpin and exhibits negligible sgRNA degradation activity, whereas rAcrIIA1 potently blocks Cas9 interference. Thus, at this locus, the RNA effector—not the protein—is the primary anti-CRISPR. This observation underscores the evolutionary plasticity of phage genomes, which can generate both RNA- and protein-based inhibitors, and suggests that redundant or compensatory mechanisms may exist to ensure robust immune evasion.

The discovery of rAcrIIA1 carries broad biological and biotechnological implications. From an evolutionary perspective, it confirms that RNA-based immune subversion is a general phenomenon across diverse CRISPR types^41^, including the prototypical Cas9 system. Phages have converged on a remarkably efficient solution: a compact ∼111-nt RNA that structurally mimics an entire sgRNA, competitively blocking native guide loading while itself remaining capable of directing Cas9 to DNA. This bifunctional nature is unique among natural anti-CRISPRs, which are typically limited to inhibition.

Our demonstration that rAcrIIA1 can be reprogrammed with arbitrary guide sequences and directs Cas9 for sequence-specific DNA cleavage *in vitro* and *in vivo* positions it as a versatile biotechnological tool (Figure 6). Importantly, rAcrIIA1 outperformed the canonical sgRNA in activating Cas9 *trans*-cleavage, suggesting its potential as an improved scaffold for molecular diagnostics^50^. In cellular assays, rAcrIIA1 guides effectively silenced an *mCherry* reporter and inhibited plasmid propagation with efficiency comparable to sgRNA, validating its utility for gene silencing and genome editing. Engineering of rAcrIIA1 may yield optimized guide RNAs with enhanced specificity, activity, or conditional control. Collectively, rAcrIIA1 not only reveals a new chapter in phage–host arms races but also expands the CRISPR-Cas9 toolbox for applications ranging from gene editing to nucleic acid detection.

## Methods

### Plasmid construction and preparation

All genetic constructs for expression were synthesized by GenScript and cloned into the modified pET28a vector with an N-terminal His_6_SUMO-tag, followed by an ubiquitin-like protein1 (Ulp1) protease cleavage site.

For crRNA and tracrRNA directed-plasmid targeting assays, SpyCas9 gene, tracrRNA and CRISPR array targeting the ampicillin resistance gene and the pMB1 ori sequence were inserted into a modified pSC101 vector. The SpyCas9 and CRISPR array were induced by l-arabinose and tetracycline, respectively. For sgRNA directed-plasmid targeting assays, Cas9 gene and sgRNA inserted into a modified pSC101 vector. The Cas9 were induced by l-arabinose. In all experiments, pUC19 was used as the target plasmid. All Acr constructs were cloned into kanamycin-resistant shuttle vector p15a, which contains T5 promoter.

For plasmid interference assays, SpyCas9 gene and rAcrIIA1 targeting the mcherry gene or the pBAD promoter were inserted into pET28a vector with an N-terminal His_6_-tag and pUC19 vector with pJ23119 promoter, respectively. Ptarget_mcherry plasmid was cloned into pET23a vector induced by l-arabinose and conferred chloramphenicol resistance.

### Protein expression and purification

All overexpressed recombinant plasmids were transformed into Rosetta (DE3) (Novagen) cells. Then all were grown to an optical density at 600 nm of 0.6–0.8 at 37 °C and induced with 0.1 mM isopropyl-1-thio-β-d-galactopyranoside (IPTG) for 14 h at 16 °C.

For SpyCas9, collected cell was resuspended in buffer A (20 mM Tris-HCl (pH 7.5) and 500 mM NaCl). After sonication and centrifugation, the supernatant was purified using Ni Sepharose (GE Healthcare), and the target proteins were eluted with buffer B (20 mM Tris-HCl (pH 7.5), 200 mM NaCl and 300 mM imidazole). Eluted protein was digested with homemade Ulp1 protease at 4 °C for 2 h and applied to a Heparin HP column (Cytiva) with elution buffer containing 20 mM Tris-HCl (pH 7.5) and 1 M NaCl. The protein was further purified by a Superdex200 Increase 10/300 gel filtration column (Cytiva) pre-equilibrated with buffer C containing 20 mM Tris-HCl (pH 7.5), 500 mM NaCl and 2 mM dithiothreitol (DTT). For AcrIIA18, harvested cell was lysed by sonication in buffer A. After centrifugation, the supernatant was incubated with Ni Sepharose (GE Healthcare). The bound protein was eluted with a buffer consisting of 20 mM Tris-HCl (pH 7.5), 500 mM NaCl, and 300 mM imidazole, and then dialyzed for 3 hours against 20 mM Tris-HCl (pH 7.5) with 150 mM NaCl, in the presence of Ulp1 protease to remove the N-terminal His6-SUMO tag. The cleaved tag was subsequently removed by incubation with Ni-NTA beads, and the protein was further purified with a HiTrap Q column (Cytiva). The protein sample was concentrated and loaded onto the Superdex200 Increase 10/300 gel filtration column (Cytiva) equilibrated with the gelfiltration buffer (20 mM Tris-HCl (pH 7.5), 150 mM NaCl, 2mM DTT). SDS-PAGE was used to analyze thepurity of protein samples in all steps.

To analyze RNA related to Cas9, pET28a_SpyCas9 plasmid (encoding an N-terminal His6SUMO-tag) and pUC19_rAcrIIA1 plasmid (containing the pJ23119 promoter) were co-transformed into Rosetta (DE3) cells. Harvested cell was lysed by sonication in a buffer consisting of 20 mM Tris-HCl (pH 7.5), 200 mM NaCl. After centrifugation, the supernatant was incubated with Ni Sepharose (GE Healthcare). The bound protein was eluted with a buffer consisting of 20 mM Tris-HCl (pH 7.5), 200 mM NaCl, and 300 mM imidazole. The eluted protein was digested with homemade Ulp1 protease at 4 °C for 2 h and further purified using a Heparin HP column (Cytiva). After concentration, the sample was used for small RNA library construction and analysis.

### *In vitro* transcription and purification of sgRNAs

All sgRNAs were transcribed in vitro using home-made T7 RNA polymerase. All transcriptional templates with a T7 promoter, RNA template sequence and ribozyme sequence were synthesized by GenScript and cloned into pUC19 vectors. The template plasmids were linearized by HindIII digestion, followed by extraction using phenol–chloroform and ethanol precipitation. The transcription reaction was carried out at 37 °C for 4 h in a buffer containing 100 mM HEPES-KOH (pH 7.9), 2 mM each rNTP, 2 mM spermidine, 15–30 mM MgCl_2_, 30 mM DTT, 0.1 mg/ml home-made T7 RNA polymerase and 40 ng/µl linearized plasmid template. Synthesized RNA was separated and gel-extracted by 12% denaturing preparative PAGE and the Elutrap Electroelution System. Finally, RNA was precipitated by alcohol and resuspended with diethylpyrocarbonate-treated water for storage at -80 °C.

### Small RNA sequencing

To profile *in vivo* Acr products following SpyCas9 induction, the pET28a vector with an N-terminal His_6_-tag harboring SpyCas9 sequence and the pET23a vector harboring Acr sequence were co-transferred into *E. coli* BL21(DE3). Overnight cultures were diluted 1:100 into 50 mL LB containing 50 μg/ml kanamycin and 100 µg/ml ampicillin. When cultures reached an OD_600_ of 0.3, induced by adding 0.1 mM IPTG at 37 °C for 3 hours. Total RNA was isolated with TRIzol Reagent according to the manufacturer’s protocol and resuspended in nuclease-free water. Sequencing libraries were constructed from the RNA using the VAHTS Small RNA Library Prep Kit for Illumina V2 (Vazyme, Cat. No. NR811). The resulting libraries were size-selected with the AMPure XP system (Beckman Coulter) and sequenced on the MGI DNBSEQ-T7 platform, generating approximately 6 × 10^6^ reads per sample. Trimmed reads were mapped to the rAcrIIA1 locus using Bowtie 2 v.2.5.4. Alignments were sorted using SAMtools v.1.19.2. Coverage tracks were generated using Geneious Prime v.2025.0.2, aligned to the nucleotide positions of the rAcrIIA1 gene.

### Plasmid targeting assays *in vivo*

For inhibition experiment, the pSC101_Cas9 plasmid, p15a_Acr plasmid and pUC19_Target plasmid were co-transformed into JM109 competent cells. A single colony from the transformation was inoculated into 5 ml of LB with 100 µg/ ml ampicillin, 50 μg/ml kanamycin and 25 µg/ ml chloramphenicol and incubated at 37℃ overnight. 50 µl culture was added to 5 ml of LB with 5 mg/ml l-arabinose, 0.1 mM IPTG and 80 ng/ml tetracycline and further grown at 37℃ for 10 hours. The cells were then serially diluted, dropped onto plates and incubated at 37 °C overnight.

For targeting experiment, the pSC101_Cas9 plasmid and pUC19_Target plasmid were co-transformed into JM109 competent cells. Colonies that grew overnight were isolated and grown in 5 ml of LB at 37 °C overnight with 100 µg/ ml ampicillin and 25 µg/ ml chloramphenicol. 50 µl culture was added to 5 ml of LB with 5 mg/ml l-arabinose and further grown at 37℃ for 10 hours. The cells were then serially diluted, dropped onto plates and incubated at 37 °C overnight.

### Plasmid interference assays *in vivo*

*E. coli* DH5α (DE3) strains carrying the pET28a_SpyCas9, pUC19_rAcrIIA1 and pET23a_mcherry plasmids were inoculated from a single colony into 5 ml of LB medium containing 50 µg/ml kanamycin, 100 µg/ml ampicillin, and 25 µg/ml chloramphenicol. The cultures were grown overnight at 37℃. Subsequently, 50 μl of the culture was transferred into 5 mL of antibiotic-free LB medium containing 5 mg/ml L-arabinose and 0.1 mM IPTG, further grown at 37℃ for 10 hours. 200 μl cultures were pelleted by centrifugation, washed twice with 1 ml PBS, and finally resuspended in 1 ml PBS. 200 µl cell suspension was aliquoted into white clear-bottom 96-well plates for analysis. Fluorescence and OD_600_ were measured using a Tecan Infinite E plex plate reader. Mcherry fluorescence was measured with an excitation wavelength of 590 nm and an emission wavelength of 635 nm. RFP fluorescence was measured in the plate reader using three biological replicates per sample.

### Fluorophore-labeled *trans*-cleavage assays

Fluorophore-labeled nonspecific ssDNA was synthesized by Sangon Biotech. For sgRNA-guided Cas9 *trans*-cleavage assays, 250 nM SpysgRNA or rAcrIIA1, 250 nM target DNA and 250 nM 5’-Cy5 labeled nonspecific substrate were added into the reaction system containing 20 mM Tris-HCl (pH 7.5), 150 mM NaCl and 10 mM MgCl_2_ in order, and 250 nM Cas9 was then added to initiate the reaction. The reaction was incubated at 37 °C for the indicated lengths of time and was terminated by adding equal volumes of loading buffer (8 M urea and 0.5 M EDTA). The reaction samples were analyzed by 20% denaturing PAGE and visualized with a Chemiluminescent Imaging System (Sagecreation).

### Electrophoretic mobility shift assay (EMSA)

Fluorescently labeled crRNA (5’-FAM-UUAACGAAUUUAUGGAUAAAGUUUUAGAGCUAUGCUGUUUUG-3’) was synthesized by Sangon Biotech. SpyCas9 protein was incubated at 0.5 μM with increasing concentrations of rAcrIIA1 (0, 0.125, 0.25, 0.5, 1, 2, and 4 μM) in a buffer containing 20 mM Tris-HCl (pH 7.5), 150 mM NaCl on ice for 15 min. Then 0.5 μM 5’-FAM-crRNA and tracrRNA were added to the reaction for another 15 min. The samples were analyzed on an 8% native PAGE and visualized using a Chemiluminescent Imaging System (Sagecreation).

### *In vitro* DNA cleavage assay

Target DNA sequence containing a 5’ -NGG-3’ PAM (for SpyCas9 and FnCas9) or 5’ - NNGRRT-3’ PAM (for sauCas9) motif was cloned into the pUC19 vector. The resultant plasmids were linearized by HindIII (NEB) for the cleavage reaction. Assays were performed in a reaction cocktail containing 20 mM Tris-HCl (pH 7.5), 150 mM NaCl, 10 mM MgCl_2_, 15 ng target DNA and 30 nM rAcrIIA1s. The reaction was initiated by adding 25 nM Cas9 proteins with a final volume of 10 µl and incubated at 37 °C for 0–15 min. Ten microliters of 50 mM EDTA was added to quench the reaction at the indicated time points. The reaction samples were then mixed with 5× prestained DNA loading buffer (Generay) and separated by 1.5% agarose gel.

For Extended Data Fig. 8a, reactions contained rAcrIIA1 (1.2–60 nM) and SpyCas9 (1–50 nM) and were incubated at 37°C for 10 min. For Extended Data Fig. 8b and 8c, reactions contained rAcrIIA1 (60–120 nM) and either FnCas9 or SauCas9 (50–100 nM), with incubation at 37°C for 60 min. All reactions were quenched by adding 10 µl of 50 mM EDTA, mixed with 5× prestained DNA loading buffer (Generay), and resolved by electrophoresis on 1.5% agarose gels.

### *In vitro* sgRNA cleavage assay

Cas9 and AcrIIA18 proteins were pre-diluted by the cleavage buffer (buffer1: 20 mM Tris-HCl, pH 7.5, 500 mM NaCl, 10 mM MgCl_2_, 5% glycerol. buffer2: 20 mM Tris-HCl, pH 7.5, 200 mM NaCl, 10 mM MgCl2, 5% glycerol). 5 μM sgRNA, 10 μM Cas9 and AcrIIA18 were used for the cleavage assay. The reaction components (Cas9, AcrIIA18 and sgRNA) were mixed in three ways. The first two components were reacted at 37℃ for 15 min, followed by the addition of the third component, and the reaction was continued for another 45 min. Then the reaction was terminated by adding equal volumes of loading buffer (8 M urea and 0.5 M EDTA). The reaction samples were analyzed by 20% denaturing PAGE and visualized by staining with Toluidine Blue.

### Complex reconstitution for Cryo-EM

For SpyCas9-rAcrIIA1 complex, purified SpyCas9 protein was mixed with rAcrIIA1 at a molar ratio of 1:1.05 and incubated on ice for 30 min. The complex sample was further purified using a Superdex200 Increase 10/300 gel filtration column with buffer containing 20 mM Tris-HCl (pH 7.5), 150 mM NaCl and 2 mM DTT. SpyCas9-rAcrIIA1-target DNA-NTS complex were reconstituted at molar ratios of 1:1.05:1.2:1.2 and purified using a Superdex200 Increase 10/300 column with buffer containing 20 mM Tris-HCl (pH 7.5), 150 mM NaCl and 2 mM DTT.

### Cryo-EM sample preparation and data acquisition

Graphene-coated holey-carbon gold grids (Quantifoil Au 300 mesh, R1.2/1.3) were glow-discharged using a PELCO easiGlow (Ted Pella) prior to cryo-EM sample preparation. SpyCas9-rAcrIIA1 and SpyCas9-rAcrIIA1-DNA complex samples were generated from size-exclusion chromatography. Freshly prepared SpyCas9-rAcrIIA1 complex sample (3.5 μL aliquots at 3.5 mg/mL) or SpyCas9-rAcrIIA1-DNA complex sample (3.5 μL aliquots at 13 mg/mL) were applied to the glow-discharged grids, blotted with Whatman No.1 filter paper at a blot force of 0 and blot time of 2.5 s (SpyCas9-rAcrIIA1 complex) or 6.0 s (SpyCas9-rAcrIIA1-DNA complex) at 4°C and 100% humidity, and plunge-frozen in liquid ethane using a Vitrobot Mark IV (Thermo Fisher Scientific).

The cryo-grids were screened and data were collected on a 200 kV Glacios^TM^ 2 microscope (Thermo Fisher Scientific) equipped with a Falcon4i detector (Thermo Fisher Scientific). Movies were recorded at a nominal magnification of 190,000×, corresponding to a calibrated pixel size of 0.74 Å. Data were collected at a dose rate of 6.7 e^−^/s/pixel, with a defocus range of −0.8 to −1.2 μm. The total electron dose was 50 electrons per Å^2^ at the specimen level. All movies were recorded semi-automatically using EPU software.

### Cryo-EM data processing

The cryo-EM data were processed using the CryoSPARC software package^51^. Motion correction of the micrographs was done using the CryoSPARC “Patch motion correction (multi)” function, and the contrast transfer function parameters were estimated using the CryoSPARC “Patch CTF estimation (multi)” function. Blob-picking, 2D classification, ab-Initio reconstruction, heterogeneous refinement, homogeneous refinement, 3D classification and non-uniform refinement were sequentially done according to the CryoSPARC data processing protocol. The resolution of the map was estimated by Fourier shell correlation analysis at a correlation cutoff value of 0.143. Local Resolution Estimation yielded global map resolution. Workflow of the data processes are illustrated in the Extended Data Figs. 5 and 7.

### Model building and refinement

Model building for the SpyCas9–rAcrIIA1 binary complex was performed using the 3.03 Å cryo-EM density map (Extended Data Fig. 5). The atomic coordinates of the SpyCas9–sgRNA binary complex (PDB 4ZT0) were docked into the density with UCSF Chimera^52^, followed by iterative manual adjustment in Coot^53^. For the SpyCas9–rAcrIIA1–DNA ternary complex, the 2.78 Å map (Extended Data Fig. 7) was used. The coordinates of the SpyCas9–sgRNA–DNA ternary complex (PDB 6O0X) served as the initial template, docked and manually refined using the same procedure. Both structures subsequently underwent real-space refinement with phenix.real_space_refine^54^, applying secondary-structure and geometric restraints. Final model quality was assessed with MolProbity^55^. All cryo-EM data processing and structure refinement statistics of the final models are summarized in Extended Data Tables 1. Structure visualizations were prepared using ChimeraX^56^ and PyMOL (http://www.pymol.org/).

## Data statistical analysis

Each experiment presented was performed with independent biological replicates. The *n*, error bars and bar graph data are defined in the figure legends. Statistical analysis of specified comparisons was performed by using a two-sided, unpaired, parametric *t*-test. Statistical significance is denoted by asterisks: \**P* < 0.05, \*\**P* < 0.01, \*\*\**P* < 0.001, \*\*\*\**P* < 0.0001; NS indicates not significant.

## Reporting Summary

Further information on research design is available in the Nature Research Reporting Summary linked to this article.

## Data availability

Atomic coordinates for SpyCas9-rAcrIIA1 binary complex (PDB: 26PH), and SpyCas9-rAcrIIA1-DNA ternary complex (PDB: 26QA) have been deposited in the Protein Data Bank (PDB) database. The cyro-EM maps for SpyCas9-rAcrIIA1 binary complex (EMDB: EMD-80808), and SpyCas9-rAcrIIA1-DNA ternary complex (EMDB: EMD-80816) have been deposited in the Electron Microscopy Data Bank (EMDB) database.

## Acknowledgments

We express our gratitude to the cryo-EM facility of Peking University Health Science Center for providing support. This work was supported by grants from, the National Natural Science Foundation of China (32371346, 32301007, 324B2056, 32471256), the Fujian Provincial Natural Science Foundation of China (2024J011007, 2023J01023, and 2023J05008), the Fundamental Research Funds for the Central Universities (20720250095 and 20720240046), the Scientific Research Foundation of State Key Laboratory of Vaccines for Infectious Diseases, Xiang An Biomedicine Laboratory (2025XAKJ0103003).

## Author contributions

L.L. conceived the study. L.L., J.C. and C.Y. designed the experiments and analysed data. Y.C. expressed and purified the proteins and prepared the cryo-EM samples. P.Z., D.C., and C.G. collected cryo-EM data. P.Z., D.C., C.G, C.Y., and L.L. carried out cryo-EM data analysis and structure determination. Y.C., and J.C. carried out all of the cloning and performed the biochemical assays. J.C., Y.C., and L.L. prepared the figures; L.L. wrote the manuscript and supervised all of the research.

## Competing interests

The authors declare no competing interests.

## Notes

### Competing Interest Statement

The authors have declared no competing interest.

